# Transcytosis-mediated anterograde transport of TrkA receptors is necessary for sympathetic neuron development and function

**DOI:** 10.1101/2022.03.03.482877

**Authors:** Blaine Connor, Naoya Yamashita, Rejji Kuruvilla

## Abstract

In neurons, many membrane proteins, synthesized in cell bodies, must be efficiently delivered to axons to influence neuronal connectivity, synaptic communication, and repair. Previously, we found that axonal targeting of TrkA neurotrophin receptors in sympathetic neurons occurs via an atypical transport mechanism called transcytosis, which relies on TrkA interactions with PTP1B, a protein tyrosine phosphatase. Here, we generated TrkA^R685A^ mice, where TrkA receptor signaling is preserved, but its PTP1B-dependent transcytosis is disrupted, to show that this mode of axonal transport is essential for sympathetic neuron development and autonomic function. TrkA^R685A^ mice have decreased axonal TrkA levels *in vivo*, developmental loss of sympathetic neurons, and reduced innervation of targets. Postnatal TrkA^R685A^ mice exhibit reduced pupil size and eyelid ptosis, indicative of sympathetic dysfunction. These findings establish the necessity of transcytosis in supplying TrkA receptors to sympathetic axons and highlight the physiological relevance of this axon targeting mechanism in the nervous system.

## Introduction

The immense length of axons imposes unique challenges on neurons in that several proteins, synthesized in cell bodies, must be efficiently delivered to axon terminals. Many axonal membrane proteins with critical functions in regulating neuronal connectivity, synaptic transmission, and nerve repair, are made in cell bodies that are meters away, and need specialized mechanisms to be transported to their final destinations. Delivery of membrane proteins to axons, after biosynthesis in neuronal soma, has been proposed to occur via several modes, including direct delivery through the secretory pathway after sorting at the trans-Golgi network, non-polarized delivery to axons and dendrites followed by selective retention in axons, or transcytosis where initial delivery of newly synthesized membrane proteins to somato-dendritic compartments is followed by endocytosis and anterograde transport (Horton and Ehlers, 2003; Winckler and Mellman, 2010). However, the functional contribution of each pathway to key processes in nervous system development, neurotransmission, and regeneration, and whether one mode of axonal transport is predominant at specific stages in the life of a neuron, remain as enduring questions in neuronal cell biology.

The family of tropomyosin-related kinases (Trk) receptors provides a prominent example of membrane proteins that undergo long-distance axonal trafficking to control neuronal survival, axon growth, and synaptic transmission (Harrington and Ginty, 2013; Scott-Solomon and Kuruvilla, 2018). In sympathetic and sensory neurons, axonal TrkA receptors are internalized after binding the ligand, Nerve Growth Factor (NGF), secreted from peripheral tissues (Cosker and Segal, 2014; Scott-Solomon et al., 2021). Axon-derived TrkA receptors are then retrogradely transported long-distance to neuronal cell bodies/nuclei to exert transcriptional control of developmental programs (Cosker and Segal, 2014; Scott-Solomon et al., 2021). The cellular and molecular mechanisms underlying retrograde TrkA trafficking are intensely investigated (Barford et al., 2017; Harrington and Ginty, 2013; Scott-Solomon and Kuruvilla, 2018). However, relatively less is known about the anterograde trafficking of TrkA receptors to ensure continued responses to target-derived ligand.

Previously, we discovered that newly synthesized TrkA receptors are delivered to axons of sympathetic neurons by transcytosis (Ascano et al., 2009; Yamashita et al., 2017). Remarkably, in contrast to the constitutive secretory pathway, anterograde TrkA transcytosis is regulated by the ligand, NGF, acting on distal axons (Yamashita et al., 2017). Since NGF is a target-derived trophic factor for sympathetic neurons and is secreted in limiting amounts during development, regulated transcytosis suggests a positive feedback mechanism that serves to dynamically scale up receptor availability in axons at times of need. We uncovered the molecular underpinnings of ligand-induced transcytosis by showing that axon-derived TrkA receptors, retrogradely transported in response to NGF, are exocytosed to soma surfaces where they interact with naive resident receptors, resulting in their phosphorylation and internalization (Yamashita et al., 2017). Endocytosed TrkA receptors are then dephosphorylated by PTP1B (Yamashita et al., 2017), a protein tyrosine phosphatase anchored at the endoplasmic reticulum (ER) with its catalytic domain localized to the cytoplasmic face (Stuible and Tremblay, 2010). Inhibition of PTP1B phosphatase activity in cell bodies of compartmentalized cultures of sympathetic neurons impairs TrkA transcytosis (Yamashita et al., 2017). In mice, sympathetic neuron-specific deletion of the PTP1B gene, *PTPN1*, results in neuron loss and impaired target innervation under limiting NGF (*NGF*^*+/-*^) conditions (Yamashita et al., 2017). Together, these findings define PTP1B as a critical regulator of TrkA transcytosis, and suggest a non-canonical role for a protein tyrosine phosphatase in positively regulating neurotrophin signaling by controlling receptor localization.

PTP1B has multiple substrates, including several Receptor Tyrosine Kinases, adhesion factors, and endosomal proteins (Stuible and Tremblay, 2010). Here, we generated TrkA^R685A^ knock-in mice, where a point mutation in TrkA abolishes binding to PTP1B, to address the essential functions of PTP1B-mediated transcytosis of TrkA receptors *in vivo*. The TrkA^R685A^ mice provide a specific genetic tool to dissect the functional outcomes of PTP1B-mediated TrkA trafficking in neurons without affecting other PTP1B substrates. We show that TrkA protein levels are drastically reduced in sympathetic axons innervating final target fields but not in ganglionic cell bodies in mutant mice, suggesting that transcytosis is a major mode for supplying TrkA receptors to axons *in vivo*. TrkA^R685A^ mice exhibit sympathetic neuron loss and target innervation defects during development, as well as dysautonomia. Thus, TrkA transcytosis mediated by PTP1B is a physiologically important mode of axonal targeting, with an essential role in sympathetic nervous system development and function.

## Results

### PTP1B-dependent transcytosis is necessary for axonal delivery of TrkA *in vivo*

Trk receptors contain a conserved PTP1B substrate recognition motif (DYYR) in their tyrosine kinase activation domain (Krishnan et al., 2015; Ozek et al., 2014). Previously, we identified a point mutation (R685A) in the PTP1B recognition motif in the TrkA receptor that abolished PTP1B binding and prevented PTP1B-mediated TrkA dephosphorylation in sympathetic neurons (Yamashita et al., 2017). Notably, we found that the TrkA^R685A^ point mutation impaired anterograde TrkA receptor transcytosis in compartmentalized cultures of sympathetic neurons (Yamashita et al., 2017). Since retrograde trafficking of axonal TrkA receptors to cell bodies is critical for NGF trophic signaling (Scott-Solomon and Kuruvilla, 2018), we asked if TrkA^R685A^ receptors are capable of undergoing retrograde transport in sympathetic neurons. To visualize TrkA trafficking, we used a live-cell antibody feeding paradigm in compartmentalized cultures of sympathetic neurons (**Figure 1A**) (Ascano et al., 2009; Yamashita et al., 2017). Compartmentalized sympathetic neuron cultures were infected with adenoviral vectors expressing FLAG-tagged chimeric receptors (FLAG-TrkB:A-WT or FLAG-TrkB:A-R685A) that have the extracellular domain of TrkB and transmembrane and intracellular domains of TrkA (Yamashita et al., 2017). Sympathetic neurons do not normally express TrkB receptors, and chimeric Trk receptors respond to the TrkB ligand, Brain-Derived Neurotrophic Factor (BDNF), but retain the signaling properties of TrkA (Ascano et al., 2009). Surface chimeric receptors were live-labeled with anti-FLAG antibody exclusively in distal axon compartments. BDNF stimulation of axons (100 ng/ml, 20 min) resulted in robust internalization of FLAG antibody-bound TrkB:A-WT and FLAG-TrkB:A-R685A receptors in distal axons (**Figures 1B-F**). BDNF treatment of distal axons for a longer period (2 hr) also resulted in the retrograde accumulation of both wild-type and mutant FLAG-labeled Trk receptors in cell bodies (**Figures 1G-K**). Together, these results indicate that axonal TrkA^R685A^ receptors are able to internalize and be retrogradely trafficked in response to neurotrophin stimulation.

**Figure 1.**
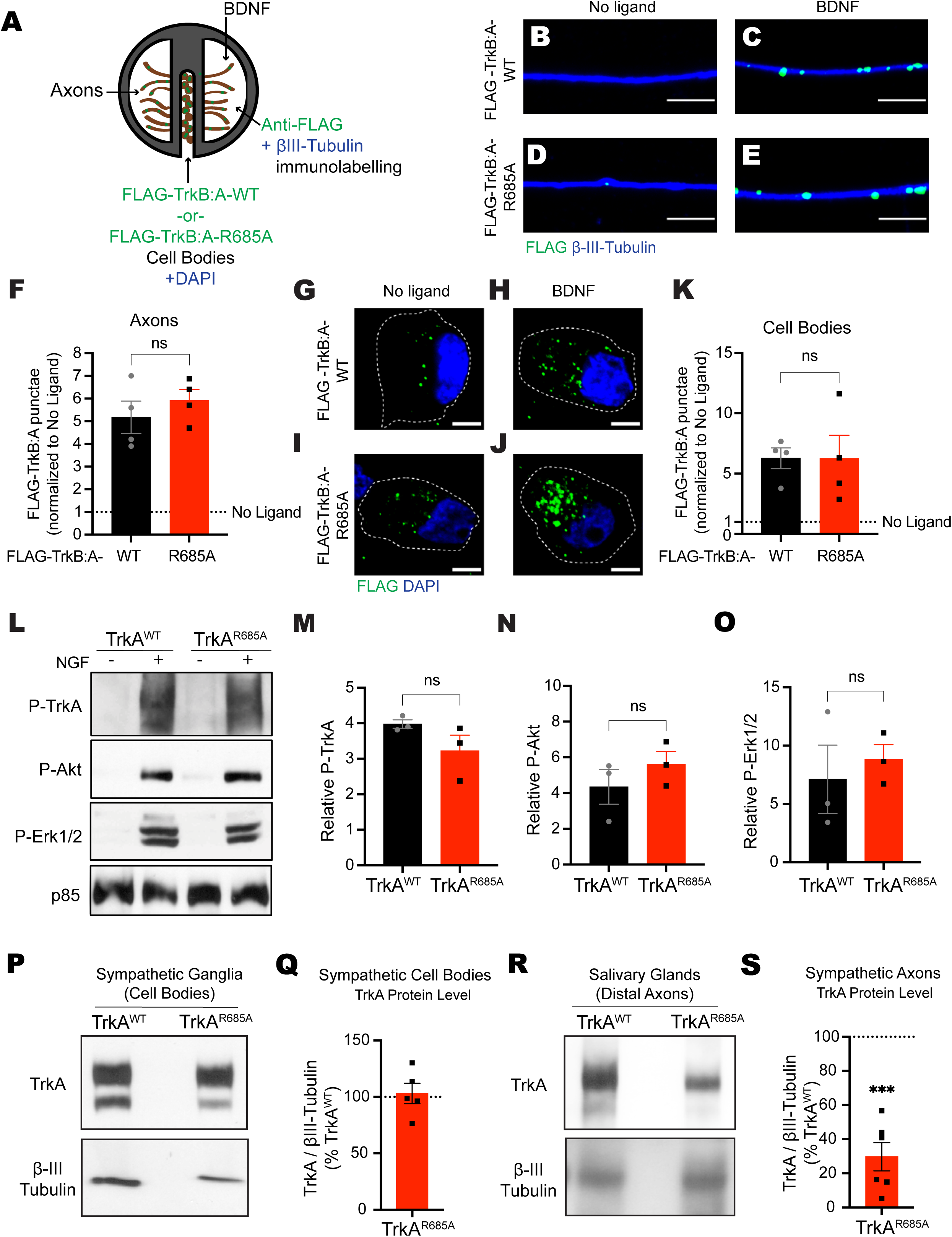
Reduced axonal TrkA levels in TrkA^R685A^ sympathetic neurons. **(A**) Antibody feeding assay to monitor retrograde trafficking of axonal TrkA receptors. Sympathetic neurons, grown in compartmentalized cultures, were infected with adenoviral vectors for FLAG-TrkB:A-WT or FLAG-TrkB:A-R685A receptors. Distal axon compartments were live-labeled with FLAG antibody, followed by stimulation with BDNF (100 ng/ml) for 20 minutes or 2 hr. **(B-E)** FLAG-TrkB:A-R685A receptors undergo ligand-induced internalization in axons, similar to FLAG-TrkB:A-WT receptors, after BDNF stimulation for 20 min. Axons were visualized by β-III-tubulin immunostaining. Scale bar: 5µm. **(F)** Quantification of FLAG-Trk punctae in axons. Results are means ± SEM from 4 independent experiments and expressed as fold-change relative to the “no ligand” condition. n.s: not significant; t-test. At least 10 axons per condition were assessed per experiment. **(G-J)** FLAG-TrkB:A-R685A receptors undergo retrograde transport to cell bodies, similar to FLAG-TrkB:A-WT receptors, after BDNF stimulation of distal axons for 2 hr. Nuclei were labeled by DAPI (blue). Scale bar: 5µm. **(K)** Quantification of FLAG-Trk punctae in axons. Results are means ± SEM from 4 independent experiments and expressed as fold-change relative to the “no ligand” condition. n.s: not significant; t-test. At least 10 cell bodies per condition were assessed per experiment. **(L)** NGF stimulation (100 ng/ml, 30 min) promotes phosphorylation of TrkA, Akt, and Erk1/2 in mass cultures of TrkA^R685A^ and TrkA^WT^ sympathetic neurons, as assessed by immunoblotting. The blots were re-probed with p85 subunit of PI-3-kinase as loading control. **(M-O)** Quantification of P-TrkA, P-Akt, and P-Erk1/2 normalized to p85 levels. Results are means ± SEM from 3 independent experiments, and expressed as fold-change relative to the “no NGF” condition for each genotype. n.s: not significant; t-test. **(P-S)** TrkA protein levels are reduced in axons innervating the salivary glands, but not in cell bodies in the superior cervical ganglia (SCG), in TrkA^R685A^ embryos. SCGs **(P)** were harvested from E16 TrkA^R685A^ mice or TrkA^WT^ littermates, and lysates immunoblotted for TrkA. Immunoblots were stripped and re-probed for β-III-tubulin as a loading control. Salivary glands lysates **(R)**, prepared from E16 TrkA^R685A^ mice or TrkA^WT^ littermates, were first subjected to TrkA immunoprecipitation and then immunoblotted for TrkA. Input lysates were immunoblotted for β-III-tubulin for protein normalization. Quantification of TrkA^R685A^ and TrkA^WT^ levels in SCGs **(Q)** and salivary glands **(S)**, were normalized to β-III-tubulin. TrkA^R685A^ expression is represented as a % of the TrkA^WT^ levels. Results are means ± SEM from 5-6 mice per genotype. ***p<0.001, t-test.

Encouraged by the results that TrkA^R685A^ mutation specifically disrupts anterograde receptor transport (Yamashita et al., 2017), without affecting retrograde trafficking, we then decided to generate TrkA^R685A^ knock-in mice to address the functional relevance of TrkA-PTP1B interactions *in vivo*. We used CRISPR/Cas9 technology to create the point mutation in the mouse *Ntrk1* gene (**Figures S1A-E**). Founder homozygous mice were back-crossed to *C57BL/6* mice for at least 3 generations. Homozygous TrkA^R685A^ mice survived to adulthood, were fertile, and had normal body weight at birth and as adults (**Figures S1F and G**). Given that the TrkA^R685A^ point mutation did not affect retrograde transport, we predicted that receptor kinase activity and downstream signaling should be preserved in the knock-in mice. To test this prediction, sympathetic neurons from neonatal TrkA^R685A^ or TrkA^WT^ mice were grown in mass cultures and stimulated with NGF (100 ng/ml) for 30 minutes. Neuronal lysates were immunoblotted for phosphorylated TrkA (P-TrkA) and canonical downstream effectors, P-Akt and P-Erk1/2. NGF stimulation increased P-TrkA, P-Akt, and P-Erk1/2 levels in TrkA^R685A^ neurons, similar to TrkA^WT^ cultures (**Figures 1L-O**). These results indicate that disruption of TrkA-PTP1B binding does not interfere with receptor kinase activity and that mutant neurons retain their ability to respond to NGF.

To ask if PTP1B-dependent transcytosis is an important means for delivering TrkA receptors to axons *in vivo*, we assessed TrkA protein levels in sympathetic nerve terminals in TrkA^R685A^ mice. We performed immunoblotting for TrkA protein in salivary glands, a target tissue that is densely innervated by sympathetic axons from the Superior Cervical Ganglia (SCG), and neuron cell bodies in the SCG in TrkA^R685A^ and wild-type littermates. We performed this analysis in mice at embryonic day 16 (E16), when sympathetic axons are beginning to reach and innervate final target organs in response to target-derived NGF (Scott-Solomon et al., 2021). Strikingly, the TrkA protein level in salivary glands of TrkA^R685A^ embryos was only ∼30% of that in wild-type animals, whereas the TrkA protein level in sympathetic ganglia was unaltered (**Figures 1P-S**). These results indicate that TrkA transcytosis is a major mode of targeting TrkA receptors to sympathetic axon terminals.

### Sympathetic neuron survival is compromised in TrkA^R685A^ mice

We next addressed the relevance of TrkA transcytosis in sympathetic neuron development *in vivo*. Tyrosine Hydroxylase (TH) immunohistochemistry and quantification of cell numbers by Nissl staining in tissue sections revealed no significant differences in the size of sympathetic ganglia (SCGs) and neuronal numbers between TrkA^R685A^ and wild-type embryos at E16 (**Figures 2A-C**). These results suggest that early developmental processes, including neuronal production, migration, and specification in the sympathetic nervous system are unaffected in TrkA^R685A^ mice (Scott-Solomon et al., 2021). However, by birth (P0), when sympathetic neurons are competing for limiting amounts of target-derived NGF for survival (Scott-Solomon et al., 2021), we found a pronounced decrease in SCG size and neuronal numbers in TrkA^R685A^ pups (**Figures 2D-F**). These phenotypes were exacerbated by 4 weeks after birth (P30) (**Figures 2G-I**). Specifically, quantification of SCG neuronal numbers revealed a substantial 41.4% decrease in P0 TrkA^R685A^ mice (12,620 ± 1500 neurons in TrkA^R685A^ mice *versus* 21,520 ± 263 in control litter-mates). The depletion of SCG neurons continued postnatally in TrkA^R685A^ mice, with the P30 mutant mice exhibiting profound neuronal loss (62% decrease) compared to wild-type litter-mates (4,750 ± 953 neurons in TrkA^R685A^ mice *versus* 12,480 ± 716 in control litter-mates). Neuronal loss in TrkA^R685A^ mice is primarily due to enhanced apoptosis, since we found a significant 3-fold increase in apoptotic cells in mutant ganglia at P0 as revealed by cleaved Caspase-3 immunostaining (**Figures 2J-L**).

**Figure 2.**
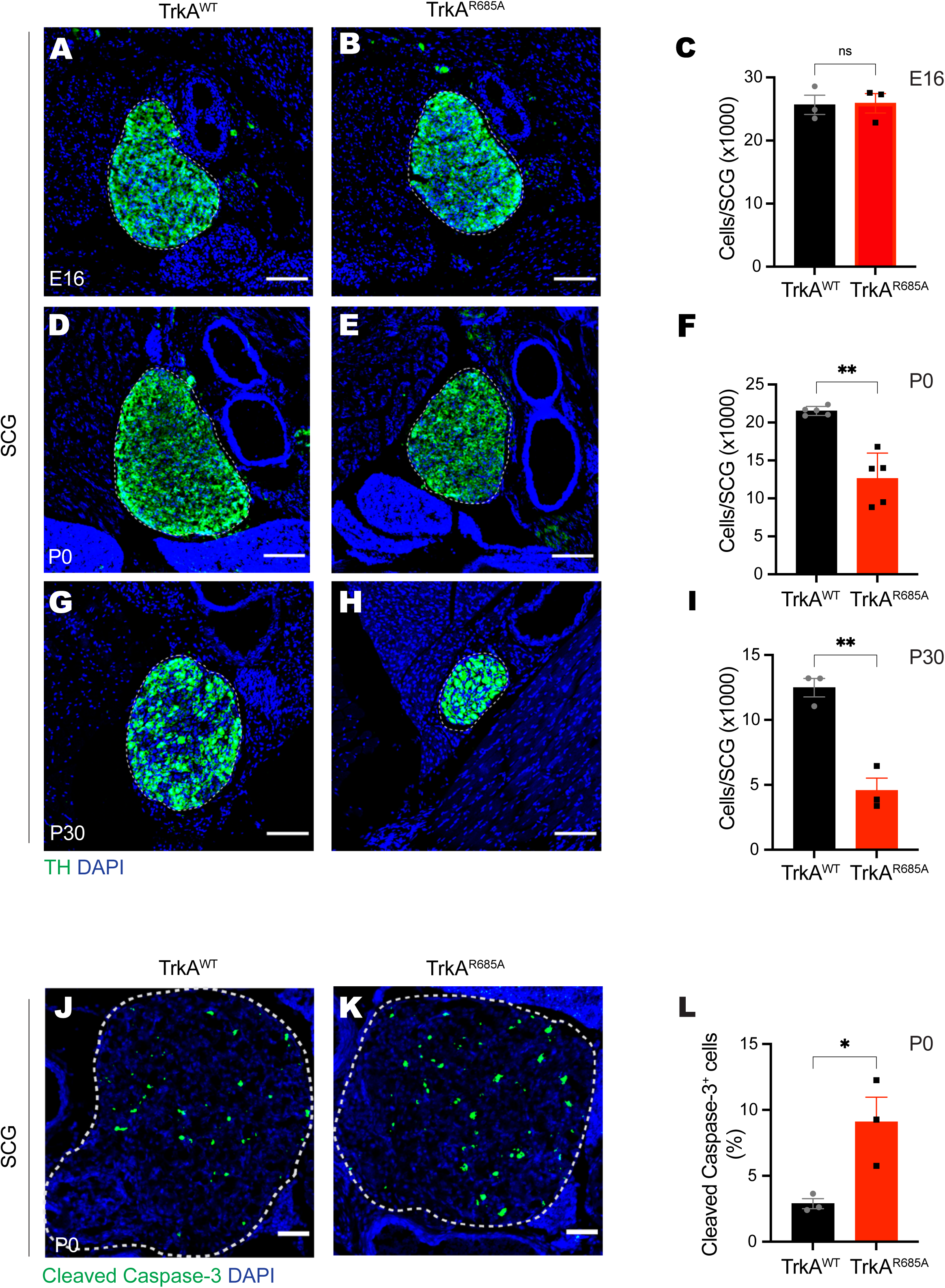
Sympathetic neuron survival is compromised in TrkA^R685A^ mice. **(A-C)** Normal SCG size and neuron numbers in E16 TrkA^R685A^ mice. **(D-I)** Decreased SCG size and neuron numbers in new-born TrkA^R685A^ mice **(D-F)**, which is exacerbated in P30 animals **(G-I)**. SCGs were visualized by TH immunostaining. Nuclei are stained with DAPI (blue). Cell counts were performed on Nissl stained SCG tissue sections. Scale bar: 100 µm. Results are means ± SEM from 3-5 mice per genotype. ns=not significant, **p<0.01, t-test. **(J-L)** Increased apoptosis in SCGs from P0 TrkA^R685A^ mice, assessed by Cleaved Caspase-3 immunostaining (green). Nuclei are stained with DAPI (blue), and SCGs are outlined by dashed line. Scale bar: 50 µm. Cleaved caspase-3-positive cells are expressed as a % of total SCG numbers. Results are means ± SEM from 3 mice per genotype. *p<0.05, t-test.

Together, these results indicate that PTP1B-mediated TrkA transcytosis is required for sympathetic neuron survival during the developmental period of NGF dependence.

### Axon innervation defects in TrkA^R685A^ mice

Given that NGF also supports sympathetic axon growth, we next assessed innervation of distal target fields using TH immunostaining. We observed decreased sympathetic innervation in several SCG targets, including salivary glands, olfactory epithelium, and the eye, in TrkA^R685A^ embryos at E16 (**Figures 3A-I**), a time when there is no neuronal loss in the ganglia (**see Figures 2A-C**). The target innervation defects persist and are aggravated, at postnatal stages in the mutant animals compared to control litter-mates (**Figures 3J-L and S2A-C**), suggesting that the reduced innervation does not reflect a developmental delay, but rather, represents the failure of sympathetic axons to grow into end-organs in response to target-derived NGF. Since TH expression is known to be regulated by TrkA signaling (Ma et al., 1992), we also assessed target innervation by immunostaining for Tuj1 (β-III-tubulin), a pan-axonal marker. We observed decreased Tuj1 immunoreactivity in E16 TrkA^R685A^ salivary glands compared to control tissues (**Figures S2D-F**), although this effect was not as severe as the decrease in TH immunoreactivity. This is likely due to sympathetic nerves constituting a fraction of total peripheral innervation.

**Figure 3.**
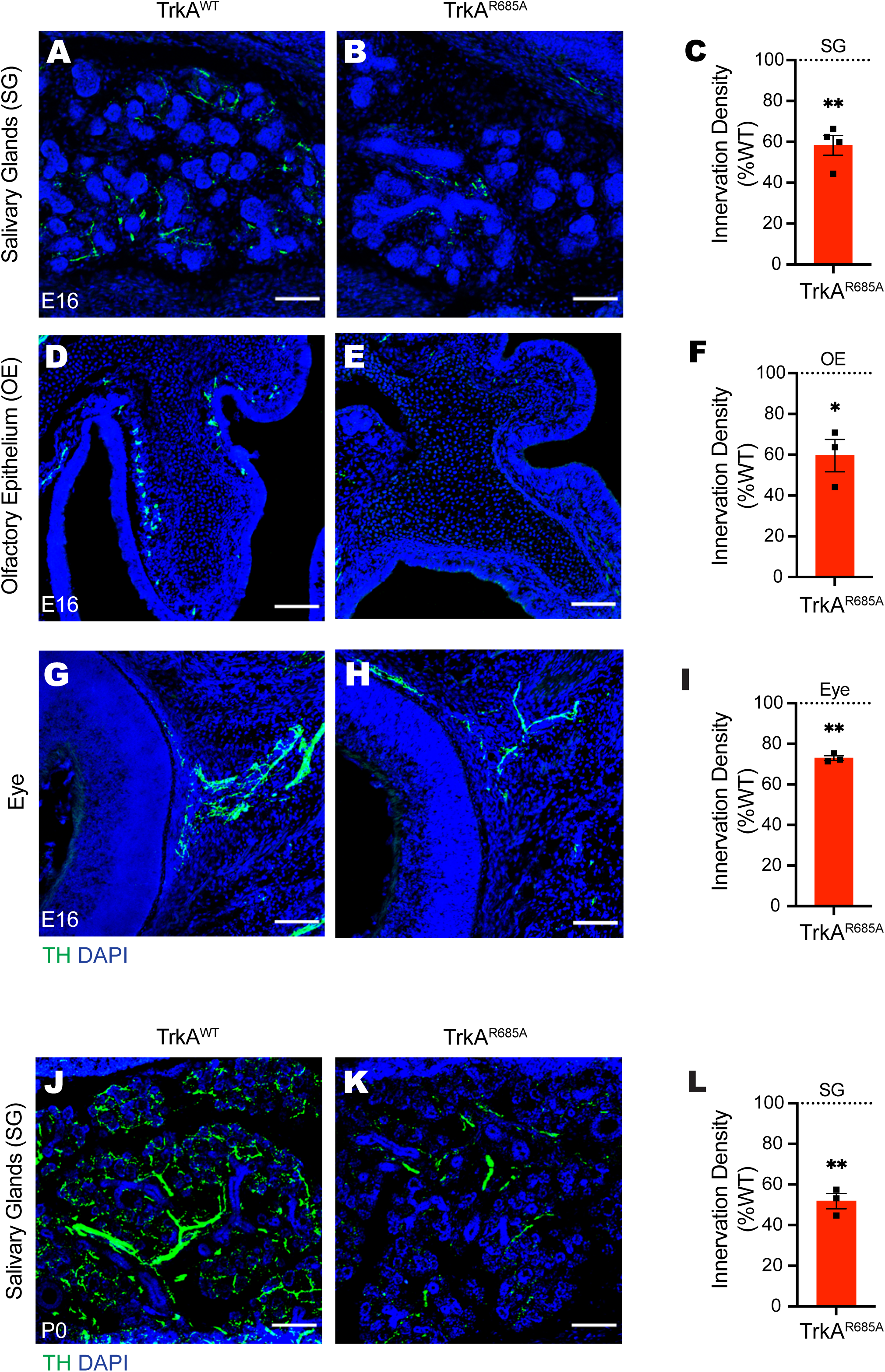
Reduced axon innervation of sympathetic targets in TrkA^R685A^ mice. **(A-I)** TH immunostaining of the salivary glands **(A-C)**, olfactory epithelium **(D-F)** and eye **(G-I)** reveals diminished sympathetic axon innervation in targets in E16 TrkA^R685A^ mice, compared to litter-mate controls. **(J-L)** Further depletion of innervation in salivary glands by birth in TrkA^R685A^ mice, revealed by TH immunostaining. Nuclei are stained with DAPI (blue). Scale bar: 100 µm. **(C, F, I, L)** Quantification of TH-positive sympathetic nerves by measuring integrated fluorescence densities per unit area using ImageJ. Results are mean ± SEM from 3-4 mice per genotype, and represented as % of TrkA^WT^ values. *p<0.05, **p<0.01, one-sample t-test with hypothetical mean of 100%.

Together, these results suggest that anterograde TrkA delivery via transcytosis is necessary for sympathetic axon growth and target innervation *in vivo*. Since the defects in target innervation precede the neuronal loss in TrkA^R685A^ mice, these results suggest that axon growth is more sensitive than neuronal survival to disruptions in axonal TrkA levels. The neuronal loss may reflect the failure of sympathetic axons to gain access to adequate levels of target-derived NGF because of the diminished innervation, as well as reduced neuronal responsiveness as axonal TrkA receptors get depleted over time.

To directly address whether the developmental phenotypes in TrkA^R685A^ mice reflected attenuated neuronal responsiveness to target-derived NGF, we tested the ability of mutant neurons to grow and survive in response to axon-applied NGF in compartmentalized cultures. NGF (30 ng/ml) added only to distal axons promotes robust growth in TrkA^WT^ neurons, with an average growth rate of ∼90 µm per day (**Figures S2G, H, and K**). In contrast, the growth-promoting effect of NGF was abolished in TrkA^R685A^ neurons (**Figures S2I, J, K**), with neurites showing retraction and degeneration. In axon growth assays, the broad-spectrum caspase inhibitor, boc-aspartyl(O-methyl)-fluoromethyl ketone (BAF, 50 µM), was added to cell bodies of compartmentalized cultures so that axon growth could be assessed independent of complications of apoptosis. We also monitored neuronal survival in response to NGF (30 ng/ml) added to distal axons in compartmentalized cultures of TrkA^R685A^ or TrkA^WT^ neurons. NGF was sufficient to support the survival of the majority of TrkA^WT^ neurons with only 9 ± 2% neurons undergoing apoptosis, assessed by Cleaved Caspase-3 immunostaining (**Figures S2L, M, and P**). In contrast, TrkA^R685A^ neurons exhibited a significant increase in neuronal apoptosis (75 ± 3.4% apoptotic neurons), despite the presence of NGF on distal axons (**Figures S2N and P**). NGF added directly to neuronal cell bodies in compartmentalized cultures is capable of promoting neuronal survival (Patel et al., 2015; Ye et al., 2003). Notably, in this condition, TrkA^R685A^ neurons were viable and healthy (8.85 ± 1.1 % apoptotic neurons) (**Figures S2O and P**), consistent with our earlier findings that the point mutation does not disrupt overall receptor signaling capacity (**see Figures 1L-O**). Together, these findings suggest that TrkA^R685A^ neurons specifically show reduced responsiveness to the NGF ligand when it is present on distal axons because of the depleted receptor levels in axons.

### Autonomic defects in TrkA^R685A^ mice

Sympathetic neurons regulate diverse physiological processes to maintain body homeostasis under basal conditions or to mobilize “fight or flight” responses to danger or stress (Scott-Solomon et al., 2021). Given the neuron loss and target innervation defects in neonatal TrkA^R685A^ mice, we asked if autonomic responses were impacted in adult mutant mice at 6-8 weeks of age. Loss of sympathetic innervation to the eye results in ptosis, or eyelid droop, in knockout mice lacking NGF or TrkA (Crowley et al., 1994; Smeyne et al., 1994), and is also observed in humans with Horner’s disease (Martin, 2018) or familial dysautonomia (Goldberg et al., 1968). Strikingly, TrkA^R685A^ mice had prominent ptosis compared to wild-type littermates (**Figures 4A-C**), suggesting that TrkA transcytosis is critical for establishing the proper sympathetic neuron connectivity necessary for autonomic functions.

**Figure 4.**
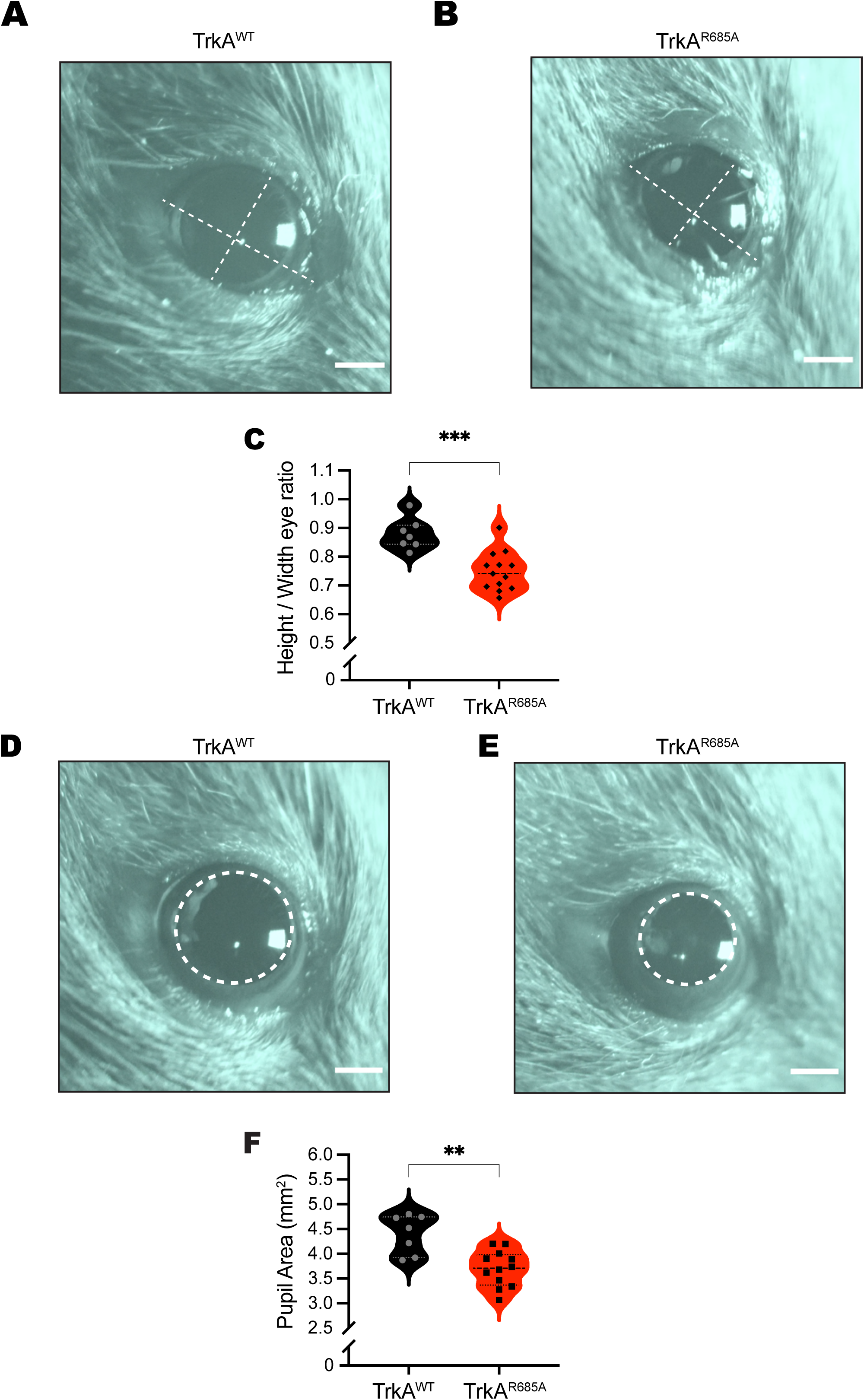
Autonomic defects in TrkA^R685A^ mice. **(A, B)** Representative images of ptosis (eyelid droop) in adult TrkA^R685A^ mice, compared to litter-mate controls. Dashed lines indicate height and width of the eye. Scale bar: 1 mm. **(C)** Quantification of ptosis (ratio of height over width of the eye). Results are the mean ± SEM from 13 TrkA^R685A^ and 7 TrkA^WT^ mice. ***p<0.001, t-test. **(D, E)** Dark-adapted TrkA^R685A^ mice have decreased basal pupil size, compared to control litter-mates. Pupil is outlined in dashed lines. Scale bar: 1 mm. **(F)** Quantification of pupil area. Results are presented as mean ± SEM from 13 TrkA^R685A^ and 7 TrkA^WT^ mice. **p<0.01, t-test.

As another assessment of sympathetic activity, we measured pupil area in mutant and control mice. In mammals, pupil size can serve as a non-invasive and rapid readout for autonomic function (McDougal and Gamlin, 2015). Pupil size is modulated by a balance of sympathetic versus parasympathetic activity, with the sympathetic component regulating pupil dilation while parasympathetic activity controls pupil constriction (McDougal and Gamlin, 2015). To measure basal pupil size, control and TrkA^R685A^ mice were dark-adapted for 2 days, and pupil sizes recorded for 30 seconds in the dark in non-anesthetized mice (Keenan et al., 2016). We found that TrkA^R685A^ mice showed a pronounced decrease in basal pupil area compared to control mice (**Figures 4D-F**). To ask whether this phenotype is due to increased parasympathetic activity, we measured pupil constriction in response to increasing light intensities, ranging from 0.01-1000 lux, administered for 30 seconds. Light onset at 0.01 lux or higher resulted in rapid constriction with greater constrictions at higher light intensities in both TrkA^R685A^ mice and control mice (**Figure S3**). The intensity response curves were virtually identical for the two groups (**Figure S3**). These results suggest that parasympathetic function is intact in TrkA^R685A^ mice, and that the smaller pupil areas likely reflect a decrease in sympathetic tone.

## Discussion

In this study, using a newly generated knock-in mouse model, we describe the *in vivo* relevance of transcytosis-mediated axonal delivery of TrkA receptors in sympathetic neuron connectivity and function. We show that a point mutation in the TrkA receptor, which disrupts TrkA transcytosis by preventing receptor interactions with PTP1B, elicits a marked depletion of TrkA protein from sympathetic axons, profound neuronal loss, reduced axon innervation of target tissues, and dysautonomia in mice. These results provide evidence that transcytosis is a key mode of targeting TrkA receptors to sympathetic axons to support neural development in response to target-derived NGF and for subsequent autonomic function.

It is striking that a point mutation in the TrkA receptor elicits robust phenotypes during sympathetic nervous system development that mimic several of the defects observed in mice with global deletion of the TrkA receptor (Fagan et al., 1996). For example, by birth, TrkA^R685A^ mice show a 41% decrease in SCG neurons, close to the magnitude of the 50% neuron loss observed in TrkA^-/-^ mice (Fagan et al., 1996). Similarly, both TrkA^R685A^ and TrkA^-/-^ mice also show reduced axon innervation of sympathetic target tissues starting at embryonic stages (∼E16.5) (Fagan et al., 1996). In contrast to the TrkA^-/-^ mice, we provide evidence, however, that the TrkA^R685A^ mutation elicits profound sympathetic abnormalities in the absence of any disruptions in receptor kinase activity, downstream signaling, or expression in ganglionic cell bodies. The most prominent cellular defect in TrkA^R685A^ neurons is the depletion of axonal TrkA receptors, which compromises neuronal responsiveness to target-derived ligand. Together, with our previous findings (Yamashita et al., 2017), we conclude that perturbed axonal targeting of TrkA receptors through transcytosis underlies the defects in sympathetic neuron development. Given that adult TrkA^R685A^ mice are viable, in contrast to TrkA^-/-^ mice, we were also able to perform physiological analyses to show that maintenance of axonal TrkA levels is critical for autonomic behaviors.

The robust sympathetic neuron phenotypes in TrkA^R685A^ mice were particularly surprising given our previous findings that conditional deletion of PTP1B did not result in overt deficits in sympathetic neuron survival or axon innervation of targets in mice (Yamashita et al., 2017). Rather, survival and innervation defects in PTP1B conditional knockout mice were only unmasked upon NGF haplo-insufficiency (*NGF*^*+/-*^) mice (Yamashita et al., 2017). One possible explanation is that since PTP1B dephosphorylates multiple substrates (Stuible and Tremblay, 2010), up-regulated tyrosine phosphorylation of one or more of these substrates could have neurotrophic effects that counter the detrimental effects of impaired TrkA targeting in PTP1B knockout mice. Thus, the TrkA^R685A^ knock-in mice provide a more precise genetic tool to dissect the interactions of PTP1B with TrkA, leaving its role in other signaling pathways intact, and affording a unique molecular handle to study the role of transcytosis in neuron development. In defining a role for PTP1B in promoting NGF-dependent trophic functions through controlling axonal localization of TrkA receptors, our findings expand the canonical view that protein tyrosine phosphatases typically diminish signaling responses to extracellular signals (Shintani and Noda, 2008; Tomita et al., 2020).

Our findings raise the possibility that different modes of axonal targeting may be utilized during distinct stages of sympathetic nervous system development. Trk receptors can be anterogradely transported to axons via the secretory pathway (Arimura et al., 2009; Tanaka et al., 2016; Vaegter et al., 2011; Zahavi et al., 2021). Therefore, constitutive delivery via the direct secretory pathway might serve to deliver TrkA receptors to growing sympathetic axons during the initial phases of axon outgrowth and extension along intermediate vascular targets, which are NGF-independent (Scott-Solomon et al., 2021). However, when sympathetic axons reach final NGF-expressing targets, PTP1B-mediated TrkA transcytosis might be a mechanism to amplify neuronal responsiveness during a developmental competition for limiting amounts of target-derived ligand. Consistent with this notion, in TrkA^R685A^ mice, we observed attenuated innervation of end-organs and loss of post-mitotic sympathetic neurons during the developmental period of known dependence on NGF trophic signaling. Together, these findings suggest that secretory transport and transcytosis likely act in a sequential manner to meet the dynamic needs of neurons during development and maturation.

The TrkA^R685A^ mice provide a powerful tool to address whether receptor transcytosis is physiologically relevant in other TrkA-expressing neuronal populations, specifically, nociceptive sensory neurons in the peripheral nervous system and basal forebrain cholinergic neurons in the brain that mediate memory and attention. In particular, basal forebrain cholinergic neurons are known to be particularly susceptible to degeneration in Alzheimer’s disease (Sofroniew et al., 2001; Whitehouse et al., 1982). It would be of interest in future studies to assess the role of aberrant TrkA receptor transcytosis in disease pathogenesis.

## Acknowledgements

We thank Haiqing Zhao and Guillermo Moya for helpful comments on the manuscript. We also thank Haiqing Zhao for guidance in the design and generation of the TrkA^R685A^ knock-in mice. We thank Aurelia Mapps and Emily Scott-Solomon for consultations on pupil analyses, and neuron survival/growth assays. We thank the JHU Transgenic Core Laboratory for help with generating the TrkA^R685A^ knock-in mice, and the JHU Integrated Imaging Center for assistance with microscopy. This work was supported by NIH R01 awards, NS114478 and NS107342, to R.K., and NIH award F31 fellowship, NS113480, to B.C.

## Author Contributions

B.C., and R.K. designed the study, with input from N.Y. during initial stages of the study. N.Y. designed and generated adenoviral vectors used in the study. B.C. conducted the experiments and analyzed data. B.C. and R.K. wrote the manuscript.

## Figure Legends

**Supplemental Figure S1.**
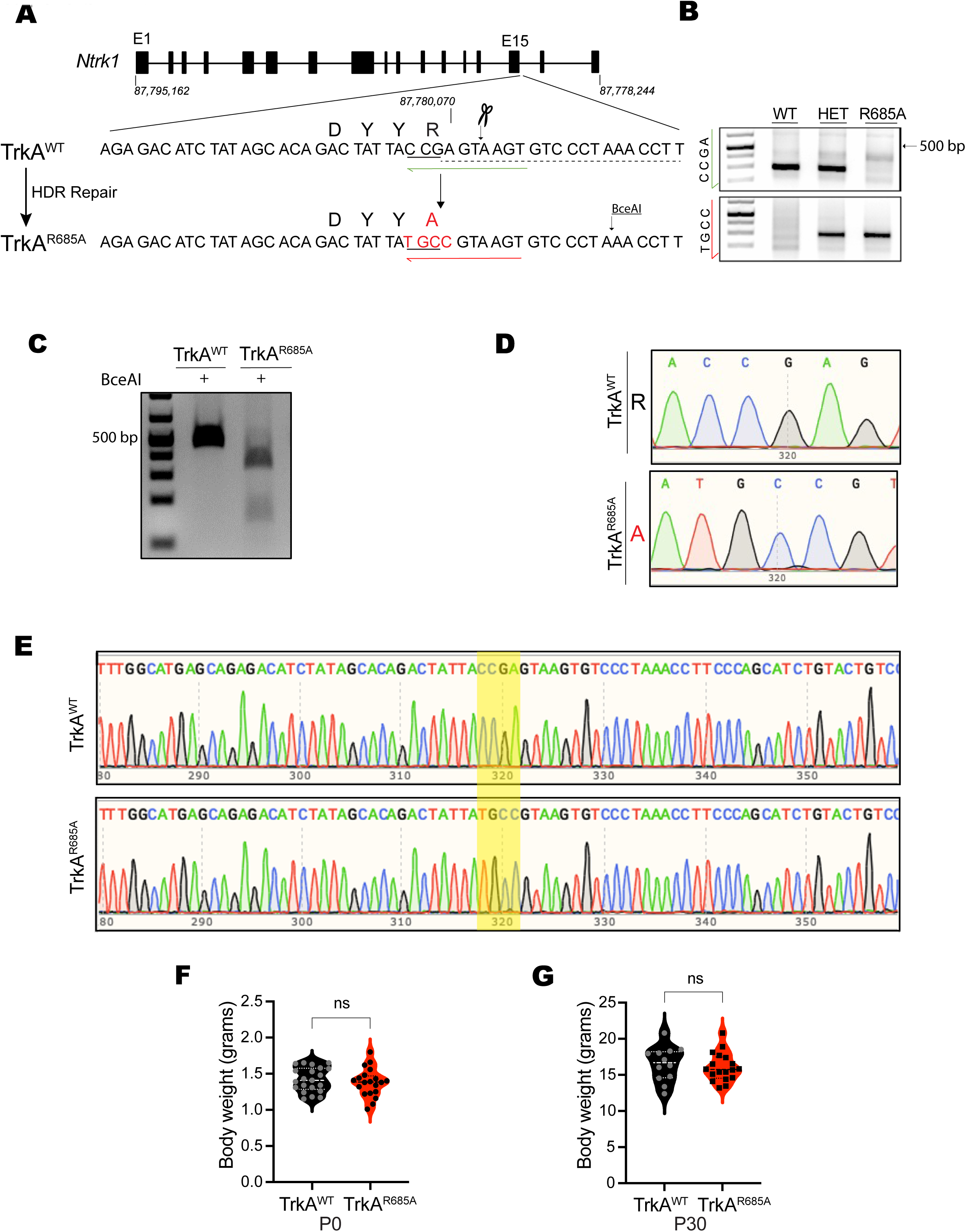
Generation of TrkA^R685A^ knock-in mice. Related to Figure 1. **(A)** Schematic for generation of TrkA^R685A^ knock-in mice. The top row shows the TrkA (*Ntrk1*) gene structure. The middle row shows the coding sequence for PTP1B interaction motif (DYYR) at exon 15 in the TrkA^WT^ allele and the Cas9 target region. Target sequence for gRNA is indicated by dashed underline, and the PAM site is indicated by solid underline. The bottom row shows the coding sequence in the TrkA^R685A^ allele after homology-directed repair (HDR). The altered nucleotide sequence, TGCC, and the corresponding amino acid, A, are shown in red. A silent C > T mutation destroys the original PAM sequence and creates a novel BceAI restriction site for Restriction fragment length polymorphism (RFLP) analysis. Green arrow (TrkA^WT^ allele) and red arrow (TrkA^R685A^ allele) indicate allele-specific primers for PCR genotyping. **(B)** PCR genotyping results. A common forward primer and allele-specific reverse primers result in a band of 287 bp for TrkA^WT^ and TrkA^R685A^. **(C)** Identification of TrkA^R685A^ knock-in animals via RFLP analysis using BceAI restriction digest. TrkA^WT^ band size: 467 bp. TrkA^R685A^ band sizes: 367bp and 100bp. **(D)** Sanger-sequencing analysis of founder mice confirms genetic modification to TrkA^R685A^. **(E)** Sequencing of TrkA^WT^ and TrkA^R685A^ alleles shows preservation of surrounding nucleotides. **(F, G)** TrkA^R685A^ animals have similar body weight as littermate controls at birth **(F)** and at postnatal day 30 **(G)**. ns=not significant, t-test.

**Supplemental Figure S2.**
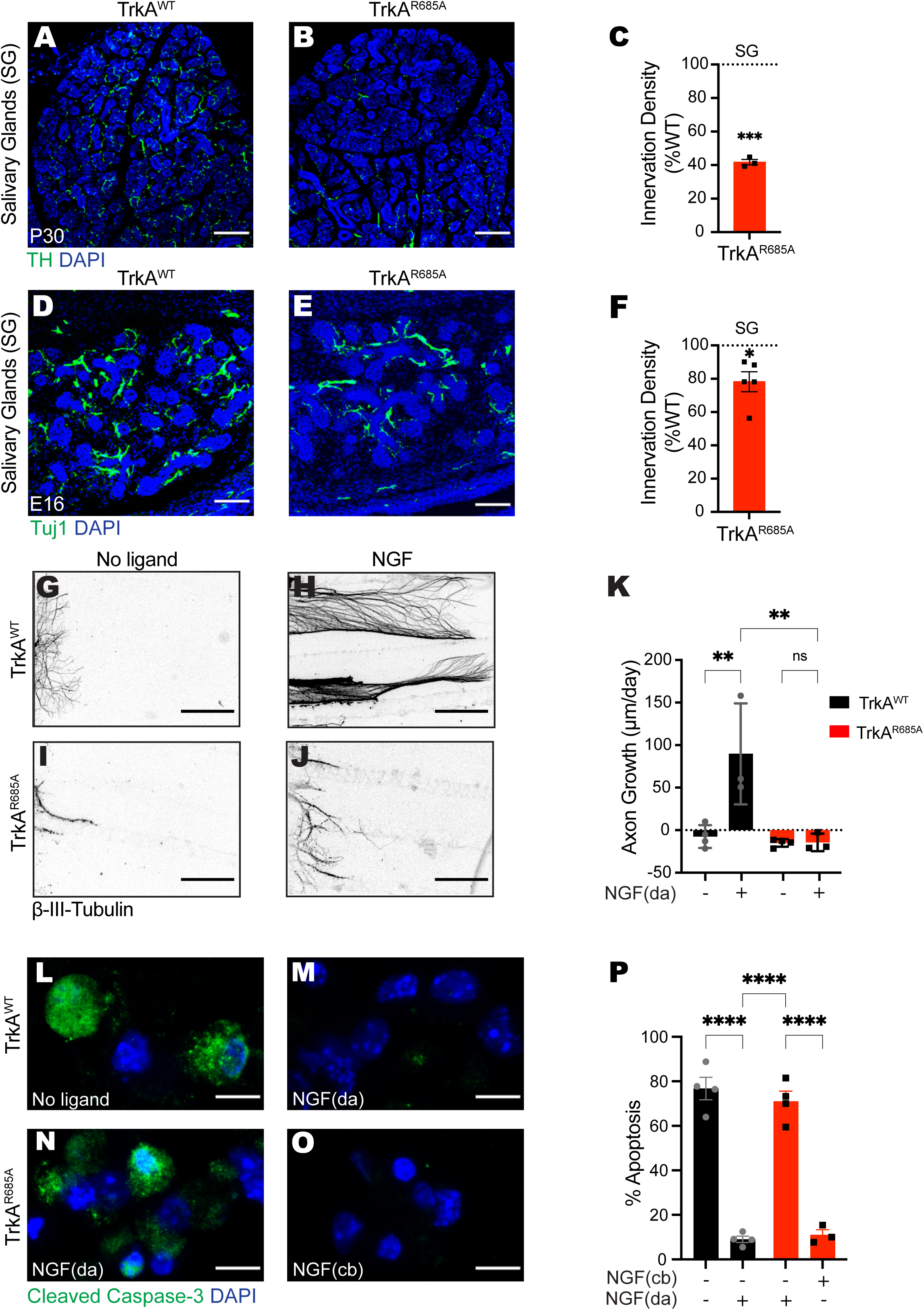
Neuron survival and growth defects in TrkA^R685A^ mice. Related to Figure 3. **(A-C)** TH immunostaining of P30 salivary glands shows marked depletion of sympathetic axon innervation in TrkA^R685A^ mice compared to litter-mate controls. **(D-F)** Immunostaining with a pan-neuronal marker, Tuj1, reveals a decrease in innervation in E16 salivary glands, compared to littermate controls. Nuclei are stained with DAPI (blue). Scale bar: 100 µm. **(C, F)** Quantification of TH-positive or Tuj1 positive nerves by measuring integrated fluorescence densities per unit area using ImageJ. Results are mean ± SEM from 3-4 mice per genotype, and represented as % of TrkA^WT^ values. *p<0.05, ***p<0.001, one-sample t-test with hypothetical mean of 100%. **(G-J)** NGF-dependent axon growth is abolished in TrkA^R685A^ neurons. Compartmentalized cultures of sympathetic neurons from P0-P3 TrkA^R685A^ or TrkA^WT^ mice were either deprived of NGF by including anti-NGF in media bathing both cell body and axon compartments **(G, I)** or maintained with NGF (30 ng/ml) added solely to distal axon compartments **(H**,**J)**. The caspase inhibitor, BAF (50 µm) was included in all analyses to prevent cell death. Panels are representative images of axons immunostained with anti-β-III-tubulin at the end of the analyses. Scale bar: 360 µm. **(K)** Quantification of rate of axon growth (µm/day assessed in 24 hr intervals for a total of 72 hr). Results are means ± SEM from 3-4 independent experiments. At least 10-15 axons per condition per experiment were analyzed. **p<0.01, two-way Anova and Tukey-Kramer post-hoc test, n.s: not significant. **(L**,**M)** NGF (30 ng/ml) added only to distal axons (da) promotes survival of TrkA^WT^ neurons, compared to the “no ligand” condition, as assessed by Cleaved Caspase-3 immunostaining. **(N**,**O)** Neuronal survival is compromised in TrkA^R685A^ neurons, when NGF (30 ng/ml) is present on distal axons (da), but not when NGF is added directly to cell bodies (cb). Neuronal nuclei were labeled with DAPI (blue). Scale bar: 5 µm. **(P)** Quantification of neuronal apoptosis by determining the percentage of neurons that were Cleaved Caspase-3-positive. Only neurons that had extended axons into side chambers, defined by the retrograde accumulation of fluorescent microspheres, were scored for apoptosis. Results are means ± SEM from 3 independent experiments. At least 20 neurons per condition were assessed per experiment. ****p<0.0001, two-way Anova and Tukey-Kramer post-hoc test.

**Supplemental Figure S3.**
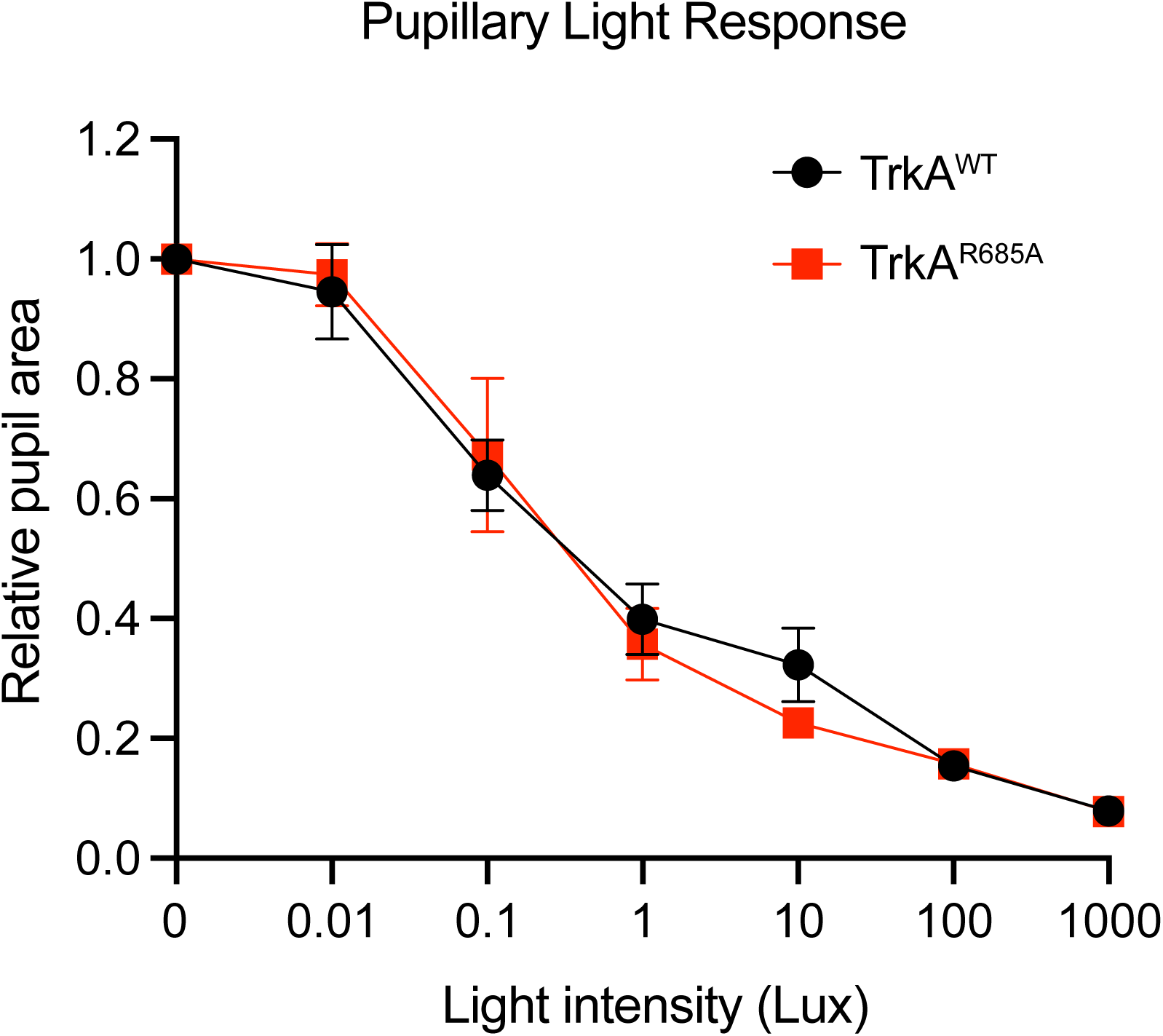
Autonomic function analysis. Related to Figure 4. Parasympathetic activity is normal in TrkA^R685A^ mice as assessed by measuring pupil constriction in response to increasing light intensities. Data are mean ± SEM from n=6 TrkA^WT^ and 5 TrkA^R685A^ animals.

## STAR METHODS

### KEY RESOURCE TABLE

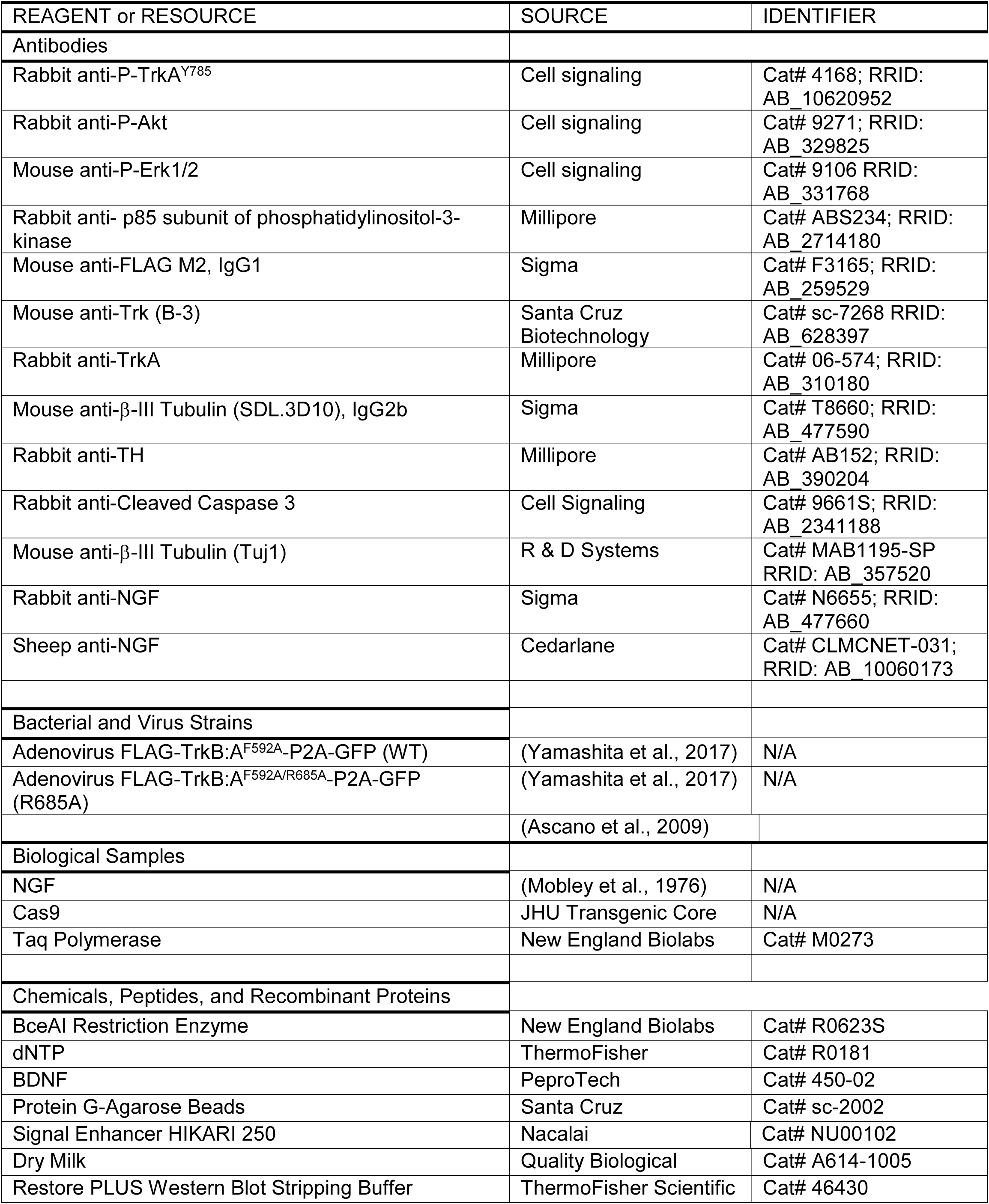
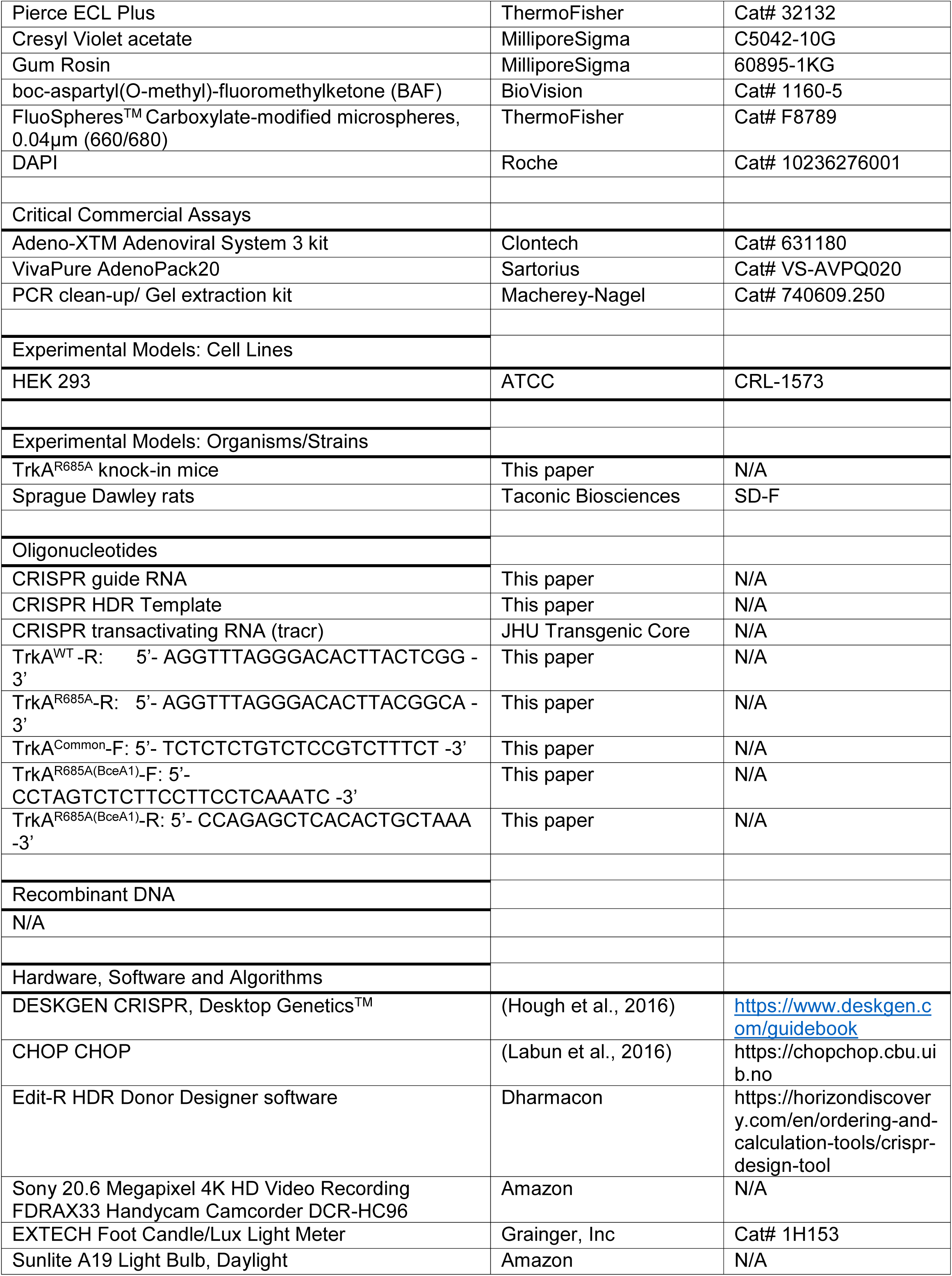
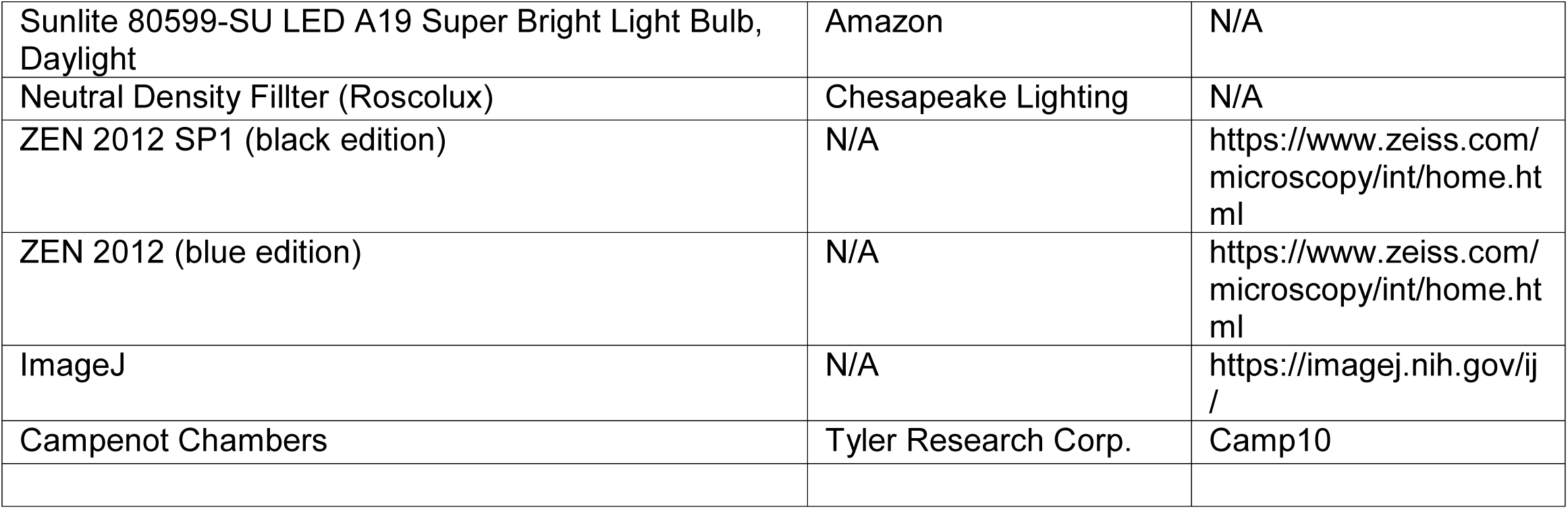

## RESOURCE AVAILABILITY

### Lead Contact

Further information and requests for resources and reagents should be directed to and will be fulfilled by the Lead Contact, Rejji Kuruvilla.

### Materials Availability

TrkA^R685A^ knock-in mice and adenoviral constructs generated in this study (see **Key Resources**) are available upon request.

### Data and Code Availability

Microscopy data reported in this paper will be shared by the lead contact upon request. This paper does not report original code.

Any additional information required to re-analyze the data reported in this paper is available from the lead contact upon request

## EXPERIMENTAL MODEL AND SUBJECT DETAILS

### Animals

All procedures relating to animal care and treatment conformed to Johns Hopkins University Animal Care and Use Committee (ACUC) and NIH guidelines. Male and female mice were included in all experiments. Mice were housed under standard conditions with access to food and water *ad* libitum. Pregnant Sprague Dawley rats were purchased from Taconic Biosciences. Dissociated cultures of sympathetic neurons were established from superior cervical ganglia (SCG) harvested from P0-P1 rat pups of both sexes as previously described (Zareen and Greene, 2009).

### Neuronal cultures

Sympathetic neurons were harvested from P0-P1 Sprague-Dawley rats or P0-P4 TrkA^R685A^ or TrkA^WT^ mice, enzymatically dissociated, and grown in mass cultures or compartmentalized cultures as described previously (Bodmer et al., 2011; Patel et al., 2015). Cells were maintained in culture with high-glucose DMEM media supplemented with 10% fetal bovine serum (FBS), penicillin/streptomycin (1U/ml), and NGF purified from mouse submaxillary glands (100 ng/ml, (Mobley et al., 1976). For NGF deprivation, neurons were placed in high-glucose DMEM containing 1.0% FBS with anti-NGF (1:1000) and boc-aspartyl(O-methyl)-fluoromethylketone (BAF) (50 μM) for 36 hr. For adenoviral infections, neuronal cultures were infected with high-titer CsCl-purified or VivaPure AdenoPack20-purified pAdenoX-Tet3G adenoviruses for 36 hr as previously described (Yamashita et al., 2017). Adenovirus-mediated protein expression was induced by adding doxycycline (Sigma, 100 ng/ml) to culture media.

## METHOD DETAILS

### Generation of TrkA^R685A^ knock-in mice

TrkA^R685A^ knock-in mice were generated using CRISPR/Cas9 technology. CHOPCHOP (Labun et al., 2016) and DESKGEN CRISPR (Hough et al., 2016) software were used to identify guide RNA target sites. *Ntrk1* sequence was obtained from the UCSC genome browser for the mm10 database (*Genome Reference Consortium m38*.*81 assembly)*. crRNA guide sequences within 50 base pairs of the desired mutation site, an on-target activity score above 45% (Doench et al., 2016), a protospacer length of 20 nucleotides and NGG protospacer-adjacent-motif (PAM) were initially selected. The sequences were scored for off-target activity (Hsu et al., 2013). The selected sequence, 5’-AAGGTTTAGGGACACTTACTCGG -3’, generated an activity score of 46% and a specificity score of 86% (Doench et al., 2016; Hsu et al., 2013). The crRNA was synthesized by Dharmacon, and the tracrRNA and Cas9 protein were provided by the Johns Hopkins University (JHU) Transgenic Core Facility. A single-stranded DNA oligonucleotide (ssODN) template with 50 base pair homology arms flanking 5’-TGCC -3’ was designed as the homology-directed repair (HDR) template. The HDR template was designed based on Ntrk1 transcript NM_001033124.1 (-) using Edit-R HDR Donor Designer software from Horizon Discovery. To prevent Cas9-mediated cleavage following the repair, the third nucleotide of the PAM recognition sequence was silently mutated (C->T), preserving the tyrosine residue in the resulting amino acid sequence and generating a novel BceAI restriction enzyme site. The HDR template was synthesized by Dharmacon, with phosphorothioate linkages between the terminal nucleotides for added stability and increased functionality. The JHU Transgenic Core performed pro-nuclear injections and implantation into C57BL/6J pseudo-pregnant females to generate TrkA^R685A^ knock-in animals. Founder mice positive for the *TrkA*^*R685A*^ allele were identified by PCR amplification, Sanger sequencing, and Restriction Fragment Length Polymorphism (RFLP) analysis using BceAI restriction enzyme, and back-crossed with C57BL6/J mice for at least 3 generations prior to experimental analyses. Colonies were maintained by mating heterozygous mice to generate knock-in homozygous mice, knock-in heterozygous mice, and littermate wild-type mice. PCR genotyping was done using a common forward TrkA primer (5’-TCTCTCTGTCTCCGTCTTTCT -3’), and reverse primer sequences for the TrkA^WT^ allele (5’-AGGTTTAGGGACACTTACTCGG - 3’) or TrkA^R685A^ allele (5’-AGGTTTAGGGACACTTACGGCA -3’). For RFLP analysis, the following primers were used; a forward primer (5’-CCTAGTCTCTTCCTTCCTCAAATC -3’) and reverse primer (5’-CCAGAGCTCACACTGCTAAA -3’). Following PCR amplification, products were subjected to BceAI digestion yielding band sizes of 467 bp for TrkA^WT^ (uncut) and 367 and 100 bp for the TrkA^R685A^ allele.

### Adenoviral constructs

Recombinant adenoviruses expressing FLAG-TrkB:A^F592A^-P2A-GFP or FLAG-TrkB:A^F592A/R685A^-P2A-GFP, referred to as FLAG-TrkB:A-WT or FLAG-TrkB:A-R685A, respectively, were previously generated by sub-cloning into pAdenoX-Tet3G using the Adeno-XTM Adenoviral System 3 kit (Yamashita et al., 2017). Recombinant adenoviral backbones were packaged into infectious adenoviral particles by transfection into HEK 293 cells using Polyethyleneimine. High-titer viral stocks were purified using a CsCl gradient or VivaPure AdenoPack20 purification kit.

### Live-cell antibody feeding

Live-cell antibody feeding assays to monitor Trk trafficking were performed as described previously (Yamashita et al., 2017). Sympathetic neurons isolated from P0-P1 rat pups were grown in Campenot chambers for 9 days in vitro (DIV) until axons projected into the side compartments. Neurons were infected with doxycycline-inducible adenoviral vectors expressing FLAG-TrkB:A-WT or FLAG-TrkB:A-R685A chimeric receptors. Neurons were treated with 200 ng/ml doxycycline for 18-24 hr to induce receptor expression. Infected neurons were identified by GFP that is co-expressed with the receptor using a self-cleaving P2A peptide sequence (GSGATNFSLLKQAGDVEENPGP). After withdrawing NGF from the culture media, surface chimeric receptors in axon compartments were labeled under live-cell conditions with a mouse anti-FLAG antibody (1:500) in Phosphate Buffered Saline (PBS) for 30 min at 4°C. Excess antibody was washed off with PBS, followed by stimulation with BDNF (100 ng/ml) added to axon compartments in culture media at 37°C for 20 min. Neurons were then placed on ice, briefly washed with ice-cold PBS, and axon compartments were washed twice in ice-cold acidic stripping buffer (0.3N Acetic Acid, 1.5M NaCl, pH 3.0) to remove surface anti-FLAG antibodies. Neurons were washed again with PBS and fixed with 4% paraformaldehyde (PFA) in PBS for 30 min at room temperature. Neurons were then permeabilized with 0.1% Triton X-100/5% normal goat serum/PBS and incubated with anti-mouse Alexa-546 secondary antibody (1:1000; Thermo Scientific) for 2 hr, then DAPI (0.3µM) for 5 min followed by mounting on slides with Fluoromount Aqueous Mounting Medium (Sigma). In some experiments, axons were also visualized by immunostaining with anti-β-III-tubulin (1:1000). To monitor retrograde trafficking of chimeric Trk receptors to cell bodies, the assays were performed as described above, except that axonal compartments were stimulated with BDNF (100 ng/ml) for 2 hr without acid stripping of surface anti-FLAG antibodies from distal axon compartments.

Images of axons and cell bodies were acquired using a LSM700 confocal microscope. The same acquisition settings were applied to all images taken from a single experiment and analyzed by using ZEN 2012 (black edition) software. Axon images were captured as a snapshot of a single plane, and somas were captured in ∼10 µm z-stacks. Intracellular accumulation of chimeric receptors was quantified as the number of FLAG-immuno-positive punctae per cell body or axon. Neurons were visualized using the GFP signal, and FLAG signals overlapping with GFP fluorescence were defined as internal receptors. For all imaging, a minimum of 10 cell bodies or axons were analyzed per condition per experiment. Results are expressed as means ± SEM and expressed relative to the “no ligand” condition.

### Immunoblotting and immunoprecipitation

To assess TrkA phosphorylation and activation of downstream signaling, sympathetic neurons isolated from P0-P3 TrkA^WT^ or TrkA^R685A^ mice were grown in mass cultures in NGF containing media for 4 days in vitro. After NGF deprivation for 24 hr in the presence of BAF (50 µM), neurons were treated with NGF (100 ng/ml) for 30 min or culture media alone (no ligand condition). Neurons were lysed in 200 μl of lysis buffer (20mM Tris-HCl (pH 8.0), 150mM NaCl, 10mM NaF, 1mM Na_3_VO_4_, 1mM EDTA, 2mm EGTA, 1% NP-40, cOmplete Mini protease inhibitor cocktail (Roche)), and total protein amounts estimated using Pierce BCA assay and a microplate reader. After normalization for protein amounts, lysates were subjected to immunoblotting with anti-P-TrkA^785^ (1:1000), followed by stripping and re-probing for P-Akt (1:1000), then P-Erk1/2 (1:1000), and lastly, p85 (1:1000) as loading control. All primary antibody incubations were done in Hikari Signal Enhancer Buffer (Nacalai) and conducted overnight for ∼16 hr, followed by 3 washes with TBST (Tris-Buffered Saline, 0.1% Tween-20) and incubations with anti-mouse- or rabbit-HRP conjugated secondary antibodies for 1 hr. Blots were then washed 3x in 5% milk/TBST and 3x in TBST prior to imaging. All immunoblots were visualized with ECL Plus Detection Reagent and scanned with a Typhoon 5 Variable Mode Imager (GE Healthcare). Results are expressed as means ± SEM and expressed relative to the “no ligand” condition.

To assess TrkA levels in sympathetic ganglia and axon terminals *in vivo*, SCG and salivary glands were dissected from TrkA^WT^ or TrkA^R685A^ E16 mouse pups. SCG lysates were subjected to immunoblotting for anti-TrkA (1:1000) in Hikari Signal Enhancer Buffer (Nacalai), then stripped with Restore PLUS Western Blot Stripping Buffer (ThermoFisher) and re-probed for anti-β-III-tubulin (1:1000). Salivary gland lysates were first subjected to TrkA immunoprecipitation using a mouse pan-Trk antibody (sc-7268). Immunoprecipitated samples were immunoblotted with rabbit anti-TrkA antibody, while input lysates were subjected to western blotting with an anti-β-III-tubulin (1:1000) antibody. Immunoblots were visualized with anti-rabbit- or -mouse-HRP conjugated secondary antibodies. All immunoblots were visualized with ECL Plus Detection Reagent and scanned with a Typhoon 5 Variable Mode Imager (GE Healthcare). Results are means ± SEM from 5-6 mice per genotype and expressed relative to TrkA^WT^ mice.

### Neuronal cell counts

Neuron counts were performed as previously described (Yamashita et al., 2017). In brief, torsos of E16, P0, or P30 mice were fixed in 4%PFA/PBS for 4 hr to overnight and cryoprotected in 30% sucrose/PBS for 24-48 hr. P30 tissues were decalcified in 0.5M EDTA prior to sucrose treatment. Torsos were then mounted in OCT and serially sectioned (12 µm). Next, every fifth section was stained with solution containing 0.5% cresyl violet (Nissl). Cells in both SCGs with characteristic neuronal morphology and visible nucleoli were counted using ImageJ.

### Immunohistochemistry

Mouse torsos were fixed in 4% PFA/PBS, and tissue sections (12 μm) were permeabilized with PBS containing 0.1% Triton X-100, and blocked using 5% goat serum in PBS + 0.1% Triton X-100. Sections were then incubated with a rabbit anti-TH antibody (1:200), mouse anti-Tuj1 antibody (1:500), or rabbit anti-cleaved caspase-3 antibody (1:200) overnight. For cleaved caspase-3, sections were first subjected to citrate antigen retrieval. Following PBS washes, sections were incubated with anti-rabbit or anti-mouse Alexa-488 secondary antibodies (1:500) and DAPI (0.3µM). Sections were then washed in PBS and mounted in Fluoromount Aqueous Mounting Medium. Images representing 6.3 μm optical slices were acquired using a Zeiss LSM 700 confocal scanning microscope with 405, 488 nm laser illumination. The same confocal acquisition settings were applied to all images taken from a single experiment. Quantification of sympathetic innervation was done by calculating integrated TH fluorescence density from multiple randomly selected images using ImageJ. Results are means ± SEM from 3-5 mice per genotype analyzed by one-sample *t*-test with hypothetical mean set at100%.

### Axon growth

Sympathetic neurons isolated from P1-P4 TrkA^WT^ or TrkA^R685A^ mice were grown in compartmentalized cultures (Campenot chambers) for 7-9 days in vitro. Neurons were either completely deprived of NGF, or NGF (30 ng/ml) was added only to distal axons. BAF (50 µM) was also included to allow assessment of axon growth without the complications of cell death. Phase contrast images of axons were captured using a Retiga EXi camera in 24-hr intervals for 3 days on a Zeiss Axiovert 200 microscope. Axon growth rate was measured using Openlab 4.0.4 for an average of 10-20 axons per condition. Axons were fixed in 4% PFA and stained with β-III-tubulin for representative images following experiments. Results are means ± SEM from 3-4 independent experiments, and analyzed using two-way Anova and Tukey-Kramer post-hoc test.

### Neuron survival

Sympathetic neurons from P1-P4 TrkA^WT^ or TrkA^R685A^ mice were grown in compartmentalized cultures (Campenot chambers) for 7-9 days in vitro to allow axon projections to distal compartments. To ensure survival scoring of only the neurons that had projected axons into the axonal compartment, fluorescent microspheres (FluoSpheres™ Carboxylate-modified microspheres, 0.04µm) were added to the distal axon compartments 24 hr before the experiments. Neurons were either starved of NGF (by adding anti-NGF at 1:1000 dilution to both cell body and distal axon compartments) or supported by NGF (30 ng/ml) added only to distal axons. After 72 hr, neurons were fixed and dying cells were visualized using cleaved Caspase-3 (1:200) immunostaining. Neuronal apoptosis was calculated by determining the percentage of neurons that had extended axons into the side chambers (visualized by fluorescent microsphere uptake) that were also positive for the cleaved Caspase-3 label.

### Pupil analyses

Pupil size measurements were performed on 6-8 week old TrkA^WT^ or TrkA^R685A^ mice as reported previously (Keenan et al., 2016). Briefly, all mice were dark-adapted and housed in single cages for 2 days and analyzed in the evenings. For all experiments, mice were un-anesthetized and restrained by hand. To mitigate stress, which can affect pupil size, researchers handled mice for several days prior to the measurements. Videos of the eye were recorded for 30 sec in the dark using a Sony 4K HD Video Recording FDRAX33 Handycam Camcorder mounted on a tripod at a fixed distance from the mouse. Manual focus was maintained on the camera to ensure that only one focal plane existed for each mouse. Pupil size was recorded under dim red light and the endogenous infrared light source of the camera to capture the basal pupil size. Results are means ± SEM from 7 TrkA^WT^ and 13 TrkA^R685A^ animals, conducted across 3 independent experiments. Student t-test was used for statistical analysis.

To assess pupillary light responses, TrkA^R685A^ or TrkA^WT^ littermates (6-8 weeks old) were individually housed in the dark as above. Light stimulus was provided by a suspended 9- or 14-watt light bulb (Sunlite A19 Light Bulb, Daylight or Sunlite 80599-SU LED A19 Super Bright Light Bulb, Daylight). Neutral density filters (Roscolux, 12.5% or 6% absorbance) were used to filter out light to estimate log unit decreases in illumination. A luminometer (EXTECH Foot Candle/Lux Light Meter, 401025) was used to set the final luminescence (lux). Un-anesthetized mice were restrained by hand in front of a mounted Sony 4K HD Video Recording FDRAX33 Handycam Camcorder in NightShot mode. Animals were held in front of the camera for 10 sec to acquire basal pupil area under infra-red light, then the light was switched on to visualize pupil constriction. Pupils were measured for 30 sec. Animals were allowed to recover for 1hr in the dark between the measurements at different light intensities. Snapshots of the captured videos were taken using QuickTime player. Pupil area at the end of each 30 sec time-point was used to assess constriction. Results are means ± SEM from 6 TrkA^WT^ and 5 TrkA^R685A^ animals, conducted across 3 independent experiments. Student t-test was used for statistical analysis.

### Ptosis

6-8 week old TrkA^WT^ or TrkA^R685A^ mice were dark-adapted and housed in single cages for 2 days. Un-anesthetized mice were restrained by hand, and images of the eye taken using a Sony 4K HD Video Recording FDRAX33 Handycam Camcorder. Images were analyzed using ImageJ to measure the width and height of the eye surface, and determine height/width ratio. Results are means ± SEM from 7 TrkA^WT^ and 13 TrkA^R685A^ animals, conducted across 3 independent experiments. Student t-test was used for statistical analysis.

## QUANTIFICATION AND STATISTICAL ANALYSIS

Sample sizes were similar to those reported in previous publications (Bodmer et al., 2011; Scott-Solomon and Kuruvilla, 2020; Yamashita et al., 2017). Data were collected randomly. For practical reasons, analyses of neuronal cell counts, axon growth, and axon innervation were done in a semi-blinded manner such that the investigator was aware of the genotypes prior to the experiment but conducted the staining and data analyses without knowing the genotypes of each sample. All Student’s t tests were performed assuming Gaussian distribution, two-tailed, unpaired, and a confidence interval of 95%. Two-way ANOVA analyses with post hoc Tukey test were performed when more than two groups were compared. Statistical analyses were based on at least 3 independent experiments and described in the figure legends. All error bars represent the standard error of the mean (SEM).

## References

Arimura, N., Kimura, T., Nakamuta, S., Taya, S., Funahashi, Y., Hattori, A., Shimada, A., Menager, C., Kawabata, S., Fujii, K., et al. (2009). Anterograde transport of TrkB in axons is mediated by direct interaction with Slp1 and Rab27. Developmental cell 16, 675–686.

Ascano, M., Richmond, A., Borden, P., and Kuruvilla, R. (2009). Axonal targeting of Trk receptors via transcytosis regulates sensitivity to neurotrophin responses. The Journal of neuroscience : the official journal of the Society for Neuroscience 29, 11674–11685.

Barford, K., Deppmann, C., and Winckler, B. (2017). The neurotrophin receptor signaling endosome: Where trafficking meets signaling. Developmental neurobiology 77, 405–418.

Bodmer, D., Ascano, M., and Kuruvilla, R. (2011). Isoform-specific dephosphorylation of dynamin1 by calcineurin couples neurotrophin receptor endocytosis to axonal growth. Neuron 70, 1085–1099.

Cosker, K.E., and Segal, R.A. (2014). Neuronal signaling through endocytosis. Cold Spring Harbor perspectives in biology 6.

Crowley, C., Spencer, S.D., Nishimura, M.C., Chen, K.S., Pitts, M.S., Armanini, M.P., Ling, L.H., MacMahon, S.B., Shelton, D.L., and Levinson, A.D. (1994). Mice lacking nerve growth factor display perinatal loss of sensory and sympathetic neurons yet develop basal forebrain cholinergic neurons. Cell 76, 1001–1011.

Doench, J.G., Fusi, N., Sullender, M., Hegde, M., Vaimberg, E.W., Donovan, K.F., Smith, I., Tothova, Z., Wilen, C., Orchard, R., et al. (2016). Optimized sgRNA design to maximize activity and minimize off-target effects of CRISPR-Cas9. Nature biotechnology 34, 184–191.

Fagan, A.M., Zhang, H., Landis, S., Smeyne, R.J., Silos-Santiago, I., and Barbacid, M. (1996). TrkA, but not TrkC, receptors are essential for survival of sympathetic neurons in vivo. The Journal of neuroscience : the official journal of the Society for Neuroscience 16, 6208–6218.

Goldberg, M.F., Payne, J.W., and Brunt, P.W. (1968). Ophthalmologic studies of familial dysautonomia. The Riley-Day syndrome. Arch Ophthalmol 80, 732–743.

Harrington, A.W., and Ginty, D.D. (2013). Long-distance retrograde neurotrophic factor signalling in neurons. Nature reviews Neuroscience 14, 177–187.

Horton, A.C., and Ehlers, M.D. (2003). Neuronal polarity and trafficking. Neuron 40, 277–295.

Hough, S.H., Ajetunmobi, A., Brody, L., Humphryes-Kirilov, N., and Perello, E. (2016). Desktop Genetics. Per Med 13, 517–521.

Hsu, P.D., Scott, D.A., Weinstein, J.A., Ran, F.A., Konermann, S., Agarwala, V., Li, Y., Fine, E.J., Wu, X., Shalem, O., et al. (2013). DNA targeting specificity of RNA-guided Cas9 nucleases. Nature biotechnology 31, 827–832.

Keenan, W.T., Rupp, A.C., Ross, R.A., Somasundaram, P., Hiriyanna, S., Wu, Z., Badea, T.C., Robinson, P.R., Lowell, B.B., and Hattar, S.S. (2016). A visual circuit uses complementary mechanisms to support transient and sustained pupil constriction. eLife 5.

Krishnan, N., Krishnan, K., Connors, C.R., Choy, M.S., Page, R., Peti, W., Van Aelst, L., Shea, S.D., and Tonks, N.K. (2015). PTP1B inhibition suggests a therapeutic strategy for Rett syndrome. The Journal of clinical investigation 125, 3163–3177.

Labun, K., Montague, T.G., Gagnon, J.A., Thyme, S.B., and Valen, E. (2016). CHOPCHOP v2: a web tool for the next generation of CRISPR genome engineering. Nucleic Acids Res 44, W272–276.

Ma, Y., Campenot, R.B., and Miller, F.D. (1992). Concentration-dependent regulation of neuronal gene expression by nerve growth factor. The Journal of cell biology 117, 135–141.

Martin, T.J. (2018). Horner Syndrome: A Clinical Review. ACS Chem Neurosci 9, 177–186.

McDougal, D.H., and Gamlin, P.D. (2015). Autonomic control of the eye. Compr Physiol 5, 439–473.

Mobley, W.C., Schenker, A., and Shooter, E.M. (1976). Characterization and isolation of proteolytically modified nerve growth factor. Biochem 15, 5543–5551.

Ozek, C., Kanoski, S.E., Zhang, Z.Y., Grill, H.J., and Bence, K.K. (2014). Protein-tyrosine phosphatase 1B (PTP1B) is a novel regulator of central brain-derived neurotrophic factor and tropomyosin receptor kinase B (TrkB) signaling. The Journal of biological chemistry 289, 31682–31692.

Patel, A., Yamashita, N., Ascano, M., Bodmer, D., Boehm, E., Bodkin-Clarke, C., Ryu, Y.K., and Kuruvilla, R. (2015). RCAN1 links impaired neurotrophin trafficking to aberrant development of the sympathetic nervous system in Down syndrome. Nature communications 6, 10119.

Scott-Solomon, E., Boehm, E., and Kuruvilla, R. (2021). The sympathetic nervous system in development and disease. Nature reviews Neuroscience 22, 685–702.

Scott-Solomon, E., and Kuruvilla, R. (2018). Mechanisms of neurotrophin trafficking via Trk receptors. Molecular and cellular neurosciences.

Scott-Solomon, E., and Kuruvilla, R. (2020). Prenylation of Axonally Translated Rac1 Controls NGF-Dependent Axon Growth. Developmental cell 53, 691–705 e697.

Shintani, T., and Noda, M. (2008). Protein tyrosine phosphatase receptor type Z dephosphorylates TrkA receptors and attenuates NGF-dependent neurite outgrowth of PC12 cells. Journal of biochemistry 144, 259–266.

Smeyne, R.J., Klein, R., Schnapp, A., Long, L.K., Bryant, S., Lewin, A., Lira, S.A., and Barbacid, M. (1994). Severe sensory and sympathetic neuropathies in mice carrying a disrupted Trk/NGF receptor gene. Nature 368, 246–249.

Sofroniew, M.V., Howe, C.L., and Mobley, W.C. (2001). Nerve growth factor signaling, neuroprotection, and neural repair. Annual review of neuroscience 24, 1217–1281.

Stuible, M., and Tremblay, M.L. (2010). In control at the ER: PTP1B and the down-regulation of RTKs by dephosphorylation and endocytosis. Trends in cell biology 20, 672–679.

Tanaka, Y., Niwa, S., Dong, M., Farkhondeh, A., Wang, L., Zhou, R., and Hirokawa, N. (2016). The Molecular Motor KIF1A Transports the TrkA Neurotrophin Receptor and Is Essential for Sensory Neuron Survival and Function. Neuron 90, 1215–1229.

Tomita, H., Cornejo, F., Aranda-Pino, B., Woodard, C.L., Rioseco, C.C., Neel, B.G., Alvarez, A.R., Kaplan, D.R., Miller, F.D., and Cancino, G.I. (2020). The Protein Tyrosine Phosphatase Receptor Delta Regulates Developmental Neurogenesis. Cell reports 30, 215–228 e215.

Vaegter, C.B., Jansen, P., Fjorback, A.W., Glerup, S., Skeldal, S., Kjolby, M., Richner, M., Erdmann, B., Nyengaard, J.R., Tessarollo, L., et al. (2011). Sortilin associates with Trk receptors to enhance anterograde transport and neurotrophin signaling. Nature neuroscience 14, 54–61.

Whitehouse, P.J., Price, D.L., Struble, R.G., Clark, A.W., Coyle, J.T., and Delon, M.R. (1982). Alzheimer’s disease and senile dementia: loss of neurons in the basal forebrain. Science 215, 1237–1239.

Winckler, B., and Mellman, I. (2010). Trafficking guidance receptors. Cold Spring Harbor perspectives in biology 2, a001826.

Yamashita, N., Joshi, R., Zhang, S., Zhang, Z.Y., and Kuruvilla, R. (2017). Phospho-Regulation of Soma-to-Axon Transcytosis of Neurotrophin Receptors. Developmental cell 42, 626–639 e625.

Ye, H., Kuruvilla, R., Zweifel, L.S., and Ginty, D.D. (2003). Evidence in support of signaling endosome-based retrograde survival of sympathetic neurons. Neuron 39, 57–68.

Zahavi, E.E., Hummel, J.J.A., Han, Y., Bar, C., Stucchi, R., Altelaar, M., and Hoogenraad, C.C. (2021). Combined kinesin-1 and kinesin-3 activity drives axonal trafficking of TrkB receptors in Rab6 carriers. Developmental cell 56, 1552–1554.

Zareen, N., and Greene, L.A. (2009). Protocol for culturing sympathetic neurons from rat superior cervical ganglia (SCG). Journal of visualized experiments : JoVE.

